# Metagenomic analysis of Pigs’ faecal microbiome and its functional response associated with dietary fibre

**DOI:** 10.1101/2023.01.29.526153

**Authors:** Keshab Barman, Manasee Choudhury, Santanu Banik, Seema Rani Pegu, Sunil Kumar, Rajib Deb, JaizIsfaqure Rahman, Swaraj Rajkhowa, Shankar Lal Jat, Sujay Rakshit, Vivek Kumar Gupta, Pranab Jyoti Das

## Abstract

Pig husbandry is the most valued and economically sustainable husbandry in the livestock farming system. The performance and productivity of the pig are chiefly dependent on the nutritional factor steered by gut beneficial bacteria. The swine gut microbiome has a direct relationship with feed efficiency. As a consequence, identifying microbial taxonomy and functional capacity is critical for proper nutrient digestion. In the present investigation, eighteen grower pigs aged 3 months and weighing 30±0.55 kg were allocated into three different groups using a randomized block layout and supplemented with QPM maize fodder of 0, 5, and 10% to the basal diet by substituting (wt/wt on DM) the maize grains, and named as T_0_, T_1_, and T_2_ to probe the increase in gut beneficial bacteria found in the faecal contents. For comparison with T_0_, T_1_, and T_2_, a random faecal sample designated R was collected from six grower pigs of the same age group and bodyweight fed a maize-soya bean-based diet. The experiment is being carried out to investigate the effect of various maize fodder levels on the metagenomic profiles of pig gut microbiota using the 16S rRNA gene. All of the experimental diets were iso-nitrogenous, with protein content ranging from 18.37 to 18.63 per cent. The data generated by 16S rRNA amplicon analysis was 68, 56, 61, and 39 Mb in R, T0, T1, and T2 samples, respectively. From taxonomic distribution, bacterial phyla namely *Firmicutes, Bacteroidetes, Proteobacteria*, and *Spirochetes* are found in descending order of relative abundance in the R, T0, and T1 groups, respectively, while *Spirochaetes, Proteobacteria, Fibrobacteres, Bacteroidetes*, and *Firmicutes*are found in descending order of relative abundance in the T2 group. The relative profusion of *Fibrobacter succinogenes* is 0.89% and 16.84% in the T1 and T2 groups. According to the findings, a higher level of maize fodder in the diet of grower pigs promotes the growth of fibre-degrading bacteria in the gut microbiota, particularly *Fibrobacter succinogenes*. Moreover, feeding green maize has decreased the population of methanobacteria in the gut of pigs, which in turn has limited the production of methane.

## INTRODUCTION

The pig is one of the world’s most lucrative farm animals used for livelihood. The performance and production of domestic animals like pig is chiefly steered by factors like genetics, environment and predominantly the diet^1^. Since pig is an omnivorous animal, diet constituents are greatly dependent not only on its digestive system but also on the microbial content of the gut. The pig gut is inhibited by a colossal miscellany of microbiota. The microbes sheltered in the gastrointestinal tract of the pig are found to have co-evolved with their hosts^2^. The architecture of the animal intestinal microbiota is shaped by the host genetic variations,^3,4^ immunity-related pathways, environmental factors such as antibiotics, social contacts and stochastic events^5,6^. Most precisely the food and predominantly energy food are decisive for the framework and activity of the gut microbiome^3,5,7–9^. In swine, the gastrointestinal chamber is the hub to trillions of bacteria, responsible for roles in host immunity, metabolism, and even behaviours to some extent^10^. In commercial pig farming, about 65-70% of the total cost of production is incurred during feed procurement. A significant way to reduce feed costs in pig husbandry is to enhance feed efficiency (FE)^11^. Carbohydrates function as the leading source of energy in pigs. In commercial pig rearing, carbohydrate solely accounts for more than 60% of the dry matter and 60-70% of the dietary energy input^12,13^. As such, the digestive enzymes produced by the animal do not quench the dietary polysaccharides aside from starch. Pigs like other mammals count on collegial gut plethora to digest this abundant feed composure and impart metabolic energy^6^. Since specific microbial taxa are in charge of the breakdown of specific carbohydrate foodstuffs, the cumulative nature of meals defines the configuration and activity of gut microorganisms^13^. Currently, the study of microbiota of animals has fascinated research as it involves the prognosis of the probable function of such diversities, which are evidenced to count on all facets of host physiology like nutrient digesting, energy utilization, and animal performance^14–17^. The latest trend in the analyses of gut microbiome mainly involves next-generation sequencing-based technology, evading the time-consuming culture-based analysis of gut diversity.

Metagenomics is currently the most preferred modus operandi to investigate the intestinal microbiota in animals, which are colonized by a complex microbial ensemble. It offers cloning and analysis of the microbial genome as entire units, without the need of culturing the individual associates. Recent research has advanced that there is a symbiotic correlation between the host and intestinal micro individuals. The sequencing of the 16S rRNA gene is a high-throughput metagenomic sequencing technology to deduce the biological benefits of microbial diversity and has been widely accepted to configure the association between the gut microbiome and host phenotypes and diseases through metagenomic wide association study^13,18^. Despite numerous approaches primarily utilizing metagenomic sequencing, a minimal is figured out about swine gut architecture^19–23^. The current study has been designed to understand the increase in gut beneficial bacteria found in the faeces of Large White Yorkshire grower pigs fed with green maize supplemented diet. The study identifies the beneficial faecal microbiota dedicated to fibre degradation as well as the abundance of bacteria present in the gastrointestinal tract of grower pigs by comparing faecal samples in the phylum, class, order, family, genus, and species hierarchy. The presumptive microorganisms identified may be interlinked with nutrient digestion and growth attributes. The present study not only characterizes the complete metagenomic profile in pigs fed with green maize but also envisages a better understanding of the digestion process in the intestine as well as provides novel information regarding the regulation of microbial diversity and its role in growth and production performance of pigs fed with green maize, thus showing a positive impact of beneficial microbe in the process of digestion.

## RESULTS

### Isolation and amplification of DNA

The DNA of all samples was isolated and the 16S rRNA gene was amplified in PCR, subsequently resolved in the agarose gel electrophoresis. The amplicon was found to be intact with the same size proving amplification of the metagenomic DNA samples.

### Faecal Microbiota DNA

The DNA of the faecal digest was isolated, fragmented, and sequenced by the MiSeq PE-150 (Illumina) platform, which generated 62 Gbp of clean reads for four samples. The average sample size was estimated at 7.4 Gbp for the high group and 8.1 Gbp for the low group.

### 16S Amplicon Analysis

All the amplicon passed through the QC has been put in the library and data were analysed using the MiSeq platform and generated quality reads for all the samples. The numbers of reads were in the range of 110,000-185,000. The data generated was found to be 68, 56, 61 and 39 MB in R, T0, T1 and T2 samples respectively. The total number of bases in R, T0, T1 and T2 was found to be 68, 574, 437, 56, 845, 541, 61, 672, 853 and 39,786, 664 respectively as shown in **Table 1**.

**Table 1:** Sample wise data statistics.

| Sr. No | Sample name | Number of Reads | Total bases | Data In Mb |
| --- | --- | --- | --- | --- |
| 1 | R | 181,122 | 68,574,437 | 68 |
| 2 | T0 | 146,670 | 56,845,541 | 56 |
| 3 | T1 | 157,420 | 61,672,853 | 61 |
| 4 | T2 | 110,667 | 39,786,664 | 39 |

### Taxonomy Annotation

The SOAP denovo software (v 1.05)^26^ was used to assemble all clean reads and mapped to microbial genomes in *National Center for Biotechnology Information* (NCBI). The aligned reads generated for each sample were used for taxonomic annotation of the microbiota present in the gut of the pigs stratified at Phylum, Class, Order, Family, Genus and Species-level and determined the abundance level. The taxonomy profile was constructed at different levels. Principal component analysis (PCA) was calculated to determine similarity and dissimilarity between the compositions of samples from the same condition. Pie charts for the sample have been plotted at six taxonomic levels. These pie charts depict the distribution of relative abundance profiles of OTUs in samples with taxonomic assignments. In the pie charts, the top nineteen categories at each of the taxonomical levels have been plotted. The category ‘Others’ includes the rest of the classifications.

### Taxonomic distribution of Random Sample

We have identified several amplicon sequence variants (ASV) whose relative abundance correlated with the feed efficiency of the diet fed to the animals. **Figure 1** represents the relative availability of microbial ASV aggregated at the Phylum, Class, Order, Family, Genus and Species level for the random feed diet of LWY grower pigs. The results revealed four beneficial bacterial phyla to be present in the majority of the faecal microbiota of the trial groups. These are-Firmicutes (31.66%), Bacteroidetes (21.11%), Proteobacteria (17.24%) and Spirochaetes (3.98%) (**Figure 1A).** At the class level, 25.25% were classified as Clostridia, 21.1% as Bacteroidia, 15.84% as Gammaproteobacteria, 4.96% as Erysipelotrichi, 3.98% as Spirochaetes and 1.45% as Bacilli (**Figure 1B**). Among other beneficial bacterial ASV detected, Epsilonproteabacteria and Deltaproteobacteria constituted 0.93% and 0.42% respectively, while 0.12% remained unassigned. The abundant orders within each bacterial community were inferred as Clostridiales, Bacteriodales, Aeromonadales, Erysipelotricales, Spirochaetales and Lactobacillales (**Figure 1C**). The most abundant family present in the random group were Succinivibrionaceae, Prevotellaceae, Ruminococcaceae and Lachnospiraceae (**Figure 1D**). At the genus level *Prevotella, Succinivibrio, Treponema, and Shuttleworthia* were classified to be present in the majority while *Roseburia, Ruminococcus* and *Campylobacter* were the least abundant (**Figure 1E**). In the species hierarchy, *Prevotellacopri* occupied 6.76% of the total, while the rest of the species remained unclassified and unassigned (**Figure 1F**).

**Figure 1:**
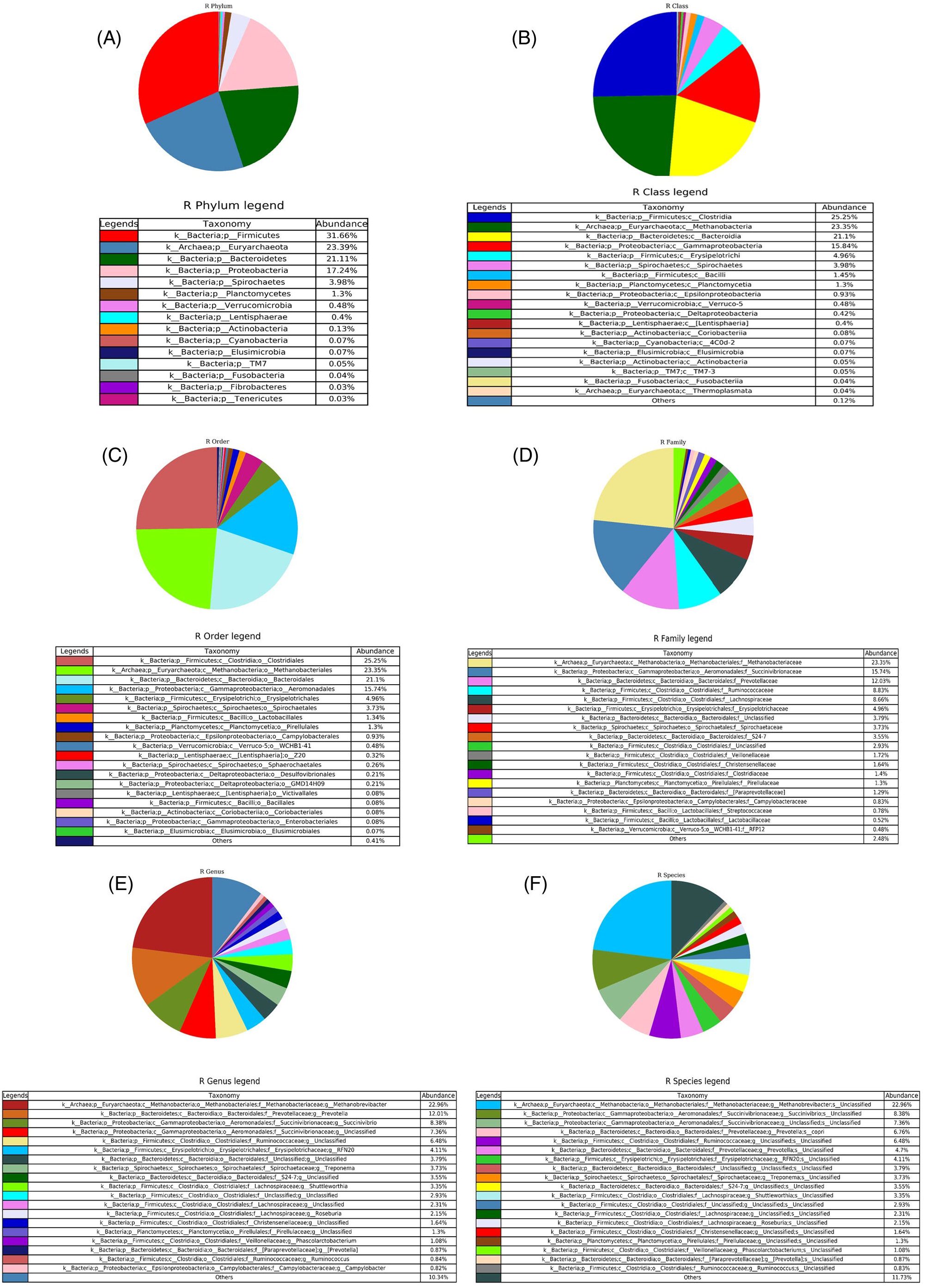
Pie chart showing the absolute abundance of each **A**. Phylum **B**. Class, **C**. Order, **D**. Family, **E**. Genus and **F**. Species in R group within each bacterial community.

### Taxonomic distribution of T0 Sample

The bacterial profusion in the faeces of pigs fed with 0% maize fodder diets was shown in **Figure 2**. The taxonomic distribution of ASV in the T0 groups showed the presence of four phyla Proteobacteria (33.82%), Bacteroidetes (22.86%), Firmicutes (18.06%) and Spirochaetes (16.37%) in abundance (**Figure 2A**). At the class level, the majority position was occupied by Epsilonproteobacteria (23.89%), Bacteroides (18.78%), Clostridia (17.16%), Spirochaetes (16.37%) and Betaproteobacteria (6.7%) (**Figure 2B**). In the order taxa Campylobacterales (23.89%), Bacteroidales (18.78%), Clostridiales (17.16%) and Sphaerochaetales (12.84%) occurred in the maximum (**Figure 2C**). From the family taxa, Campylobacteraceae (22.6%) holed a wider space followed by Sphaerochaetaceae (12.84%), Bacteroidaceae (10.51%), Porphyromonadaceae (7.1%), Ruminococcaceae (6.94%) (**Figure 2D**). Amid the genera *Arcobacter* (22.5%), *Sphaerochaeta* (12.84%) and *Bacteroides* (10.35%) were revealed to present in plenty (**Figure 2E**). The species *Arcobacter cryaerophilus* (18.26%) was detected as the most abundant of all the bacterial communities while *Wolinella succinogenes* (1.29%) occupied a minor percentage (**Figure 2F**). The rest of the species remained unclassified.

**Figure 2:**
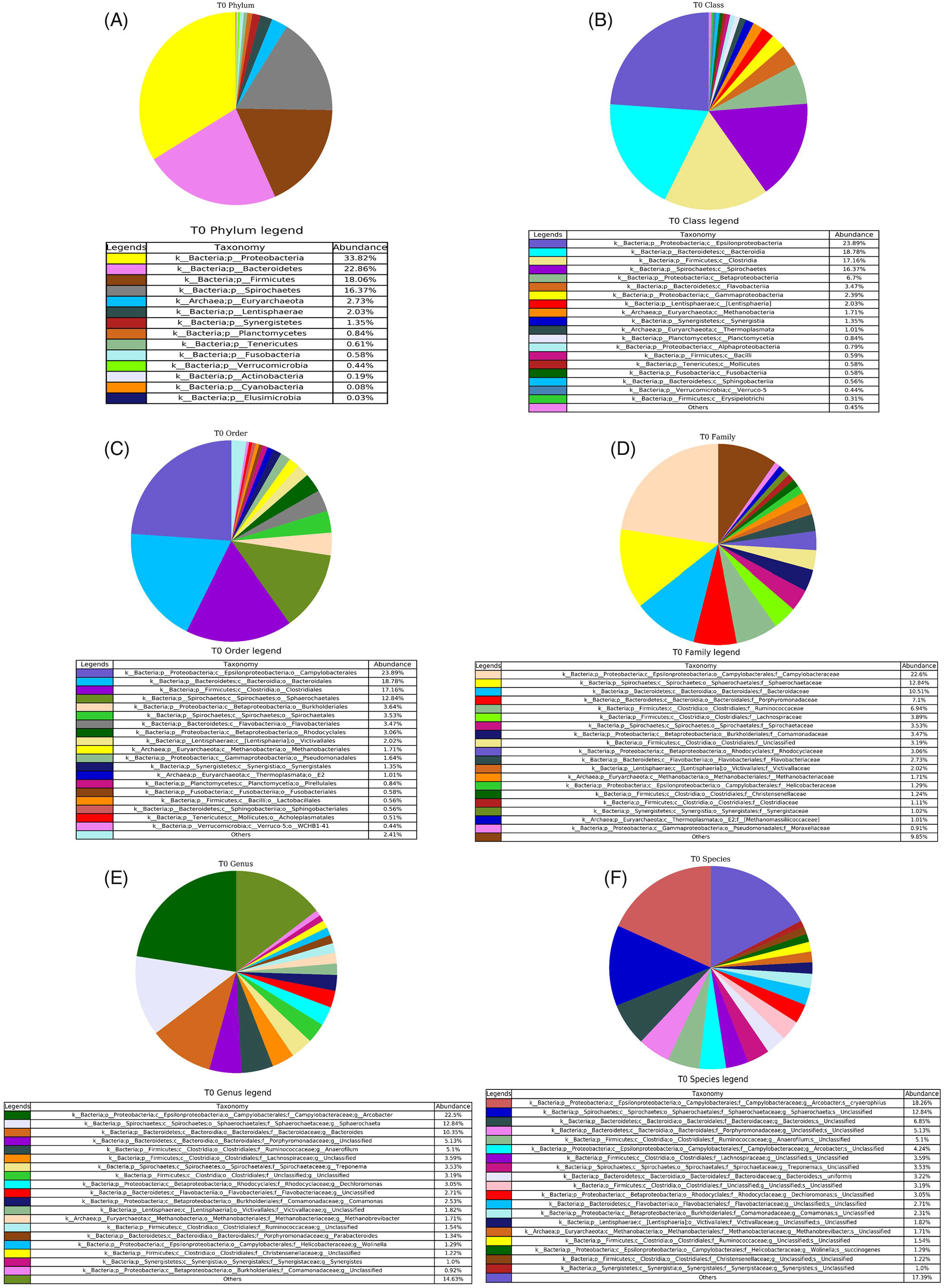
Pie chart showing the absolute abundance of each **A**. Phylum **B**. Class, **C**. Order, **D**. Family, **E**. Genus and **F**. Species in T0group within each bacterial community.

### Taxonomic distribution of T1 Sample

The taxonomic distribution of the bacterial communities in the T1 groups is shown in **Figure 3**. The gastrointestinal tract of the pigs fed with 5% maize diet contained the majority of the phylum Proteobacteria (28.01%), Spirochaetes (17.78%), Bacteroidetes (14.06%) and Firmicutes (13.98%) (**Figure 3A**). The presence of phylum Fibrobacteres (0.89%) has been detected for the first time in this study. A greater quantity of Epsilonproteobacteria (23.87%), Spirocaetes (16.82%), Bacteroidia (13.79%), Clostridia (12.17%), appeared at the class level (**Figure 3B)**. Bacteria belonging to the order Campylobacterales (23.87%), Bacteroidales (13.79%), Clostridiales (12.15%) and Spirochaetales (10.11%) absorbed most of the space (**Figure 3C)**. The abundant family was inferred to be Campylobacteraceae (23.73%), Spirochaetaceae (10.11%) and Sphaerochaetaceae (6.71%) (**Figure 3D)**. The genera included *Arcobacter, Treponema* and *Sphaerochaeta* in the largest community (**Figure 3E)**. The most plentiful species identified is *Arcobacter cryaerophilus* (23.23%) (**Figure 3F)**.

**Figure 3:**
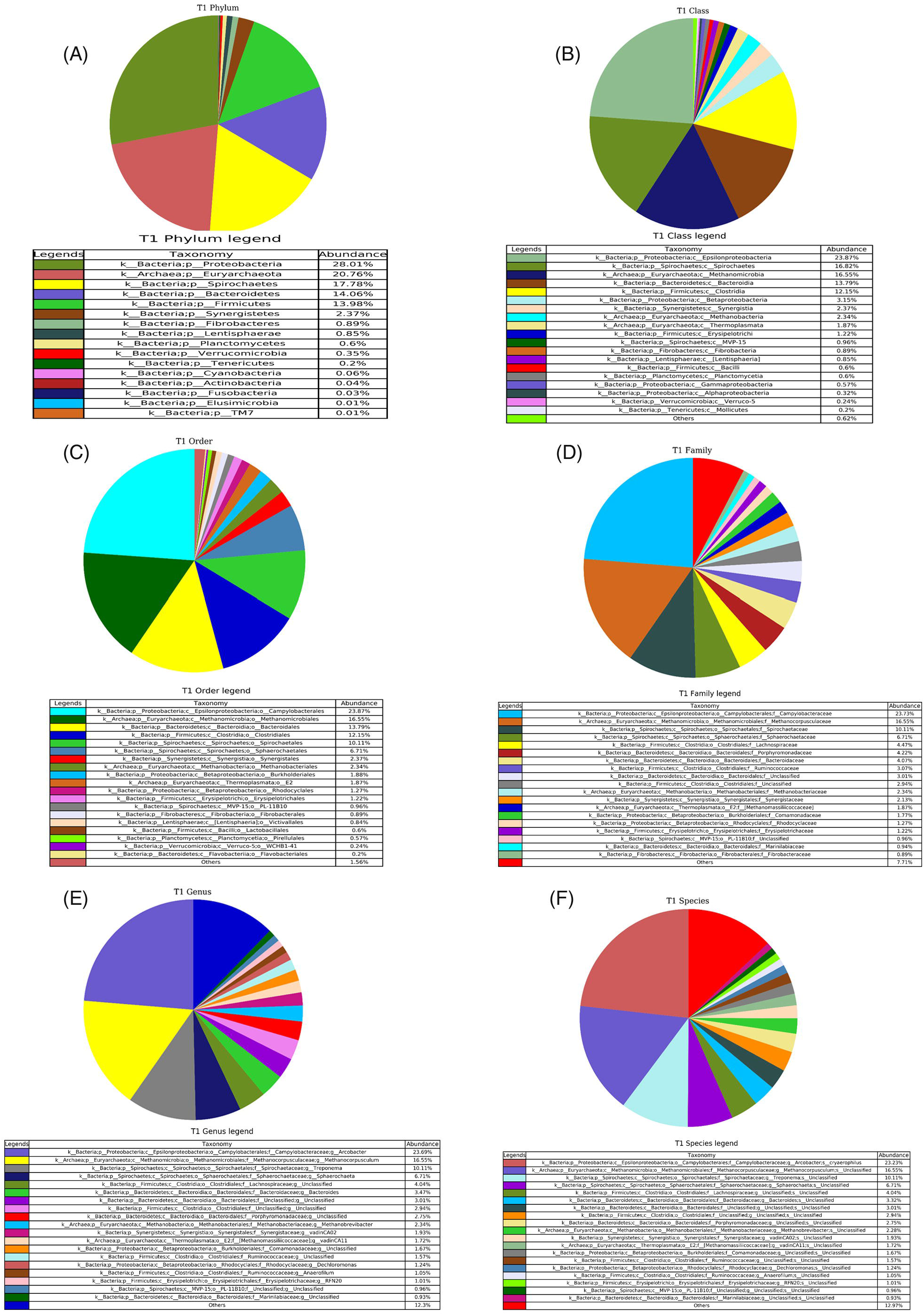
Absolute abundance were shown in Pie chart **A**. Phylum **B**. Class, **C**. Order, **D**. Family, **E**. Genus and **F**. Species in T1 group within each bacterial community

### Taxonomic distribution of Sample T2

The faecal microbiota of the pigs fed with a 10% maize diet (**Figures 4**) divulged the presence of five phyla in a large aggregate *viz*. Spirochaetes (39.08%), Proteobacteria (21.53%), Fibrobacteres (16.84%), Bacteroidetes (10.58%) and Firmicutes (7.52%) (**Figure 4A)**. Following this, the hierarchy was investigated to have a bountiful bacterial ASV belonging to the class Spirochaetes (37.94%), Epsilonproteobacteria (20.21%), Fibrobacter (16.84%), Bacteroidia (10.52%) and Clostridia (6.97%) (**Figure 4B)**. A parallel trend was observed in the order and family of the respective classes of bacteria **(Figure 4C** and **Figure 4D)**. The genus *Treponema* (37.12%), *Arcobacter* (20.09%), and *Fibrobacter* (16.84%) (**Figure 4E**) and the species *Arcobacter cryaerophilus* (19.68%) and *Fibrobacter succinogenes* (16.84%) were discovered to be present in maximum in the T2 groups (**Figure 4F**).

**Figure 4:**
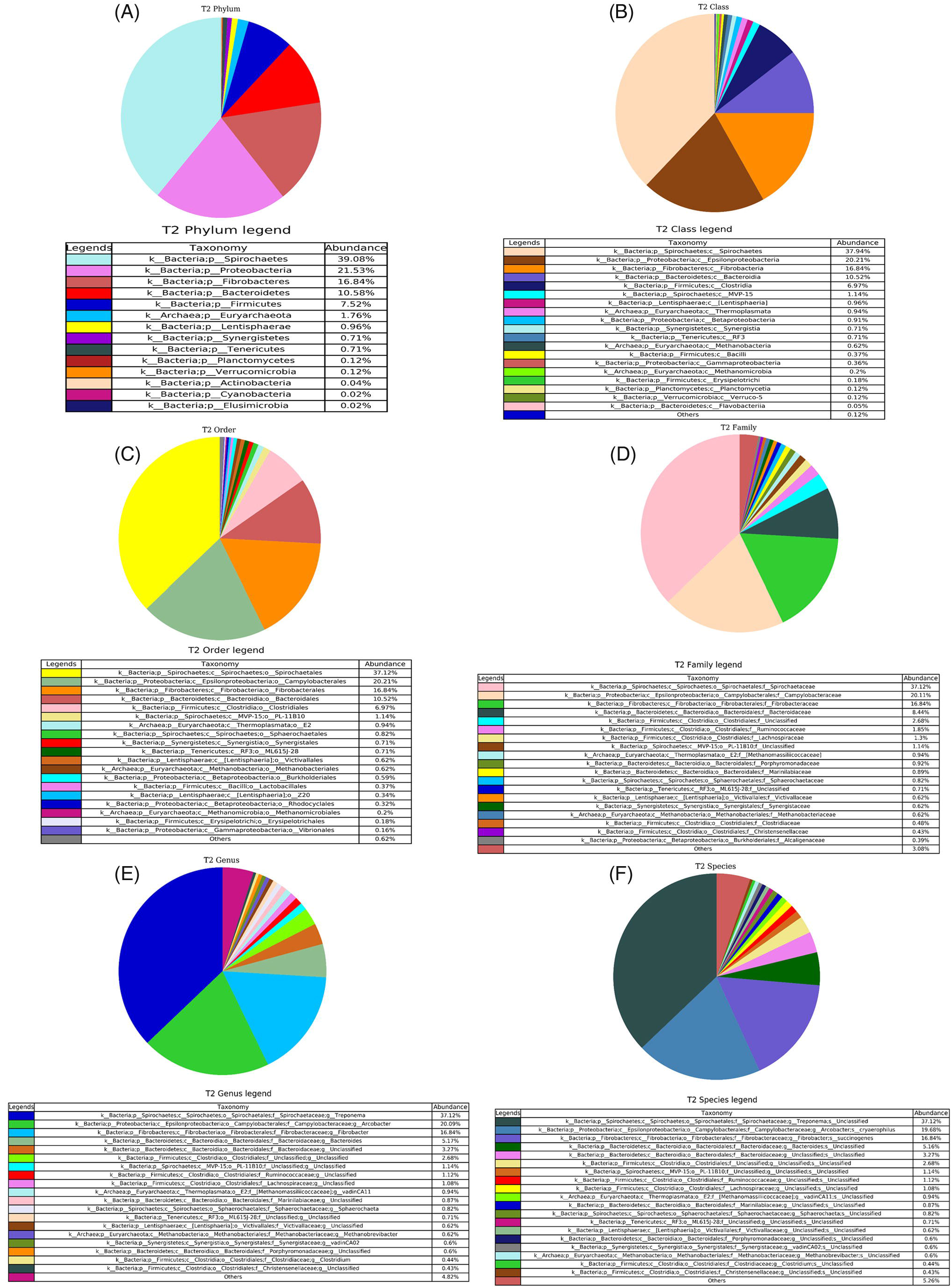
Absolute abundance were shown in Pie chart **A**. Phylum **B**. Class, **C**. Order, **D**. Family, **E**. Genus and **F**. Species in T2 group within each bacterial community

### Comparative Analysis between samples at each taxonomic level

**Figure 5A** shows the comparative analysis of the microbiotas at the phylum level. The abundance of phylum Firmicutes and Bacteroidetes slackened from the random groups to the T2 groups. A significant drop was seen in the abundance of Euryarchaeota in the T2 groups compared to the random. A spiking increase was seen in the phylum Fibrobacteres in the T2 groups (16.8%) compared to Random (0%). An increasing trend was also seen in the phylum Spirochaetes (T2-39.1%). Figure **5B**, **C** and **D**, represent the relative difference in the taxa of the experimental animals in the class, order, and family and **Figure S1 A** and **B**, represent genus and species level. Members of the class Clostridia, Methanobacteria, Bacteroidia, Gammaproteobacteria, Erysipelotrichi and Bacilli decreased from the random to T2 groups while bacteria of the class Fibrobacteria and Spirochaetes increased in their abundance with the increase in the fibre diet. Similar kinds of trajectories have been observed in the respective order, family and genus ranking of the microbiotas. The species *Fibrobacter succinogen* increased in significant abundance with the maize-fed diet (T2-16.8%) compared to the random. On the other side, no individuals of the species *Prevotellacopri, Prevotella stercorea* and *Faecalibacter iumprausnitzii*, present in the random were identified in the gut of T0, T1 and T2 animals. Species *Arcobacter cryaerophilus* is found to be present equivalently in all the trialled groups except the random. The comparative scrutiny between the samples has been done at different taxonomic levels taking a 0.5% threshold.

**Figure 5:**
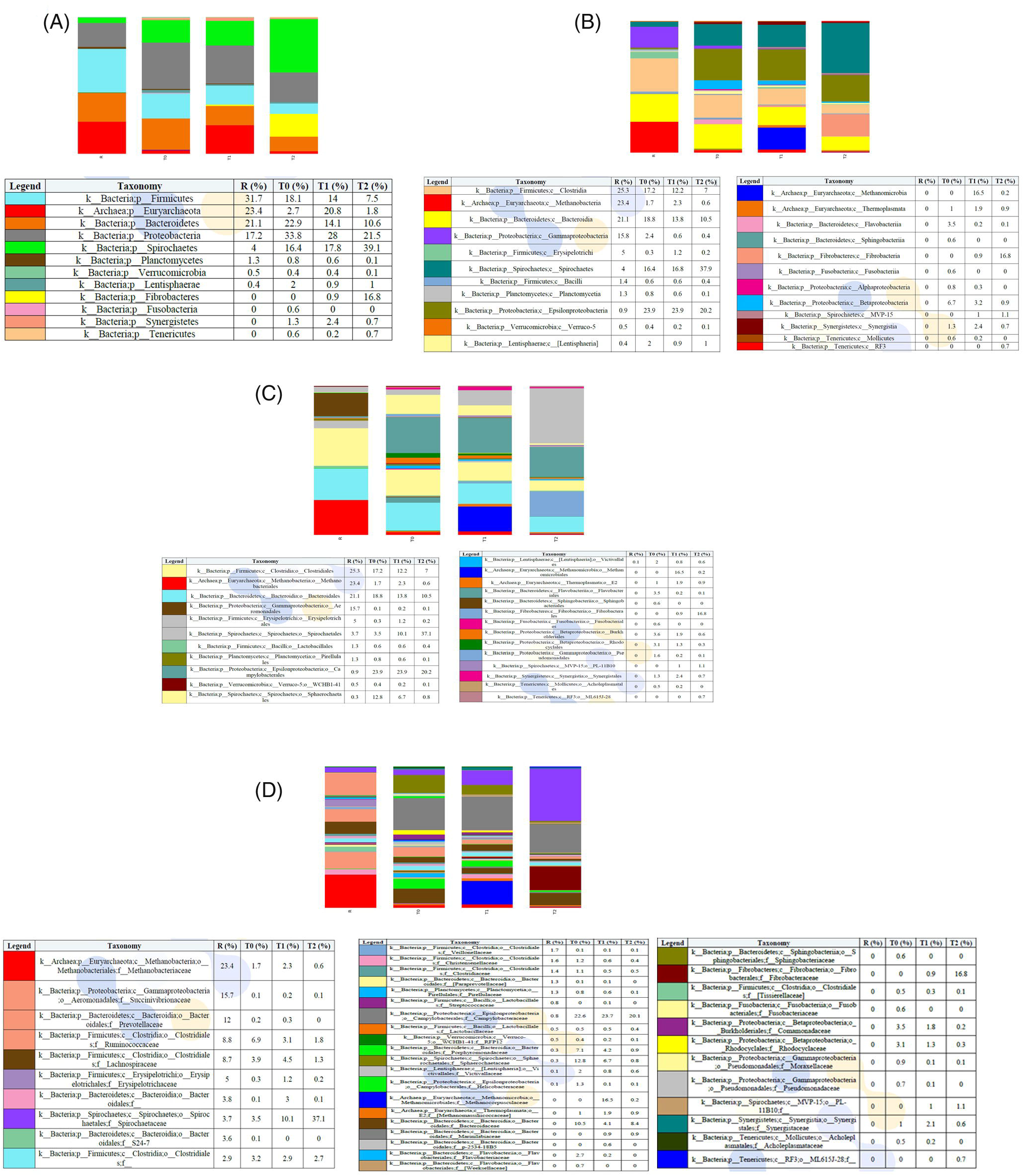
Stacked Bar chart showing the relative abundance of each **A**. Phylum **B**. Class, **C**. Order, and **D**. Family within each sample

### Alpha diversity

Alpha diversity outlines the diversity of organisms in a sample with a single number. The alpha diversity of the pig gut microbiome was measured by Shannon diversity indices. **Table 2** shows the microbial alpha diversity at the species level influenced by different feed compositions. The table interprets that there occurs a precise difference in alpha diversity of gut microbiota after feeding of different diet rations. Alpha diversity of the gut microbiota decreased synchronously with the change in dietary supplements. A total of 548, 450, 479 and 203 OTUs were detected in the random, T0, T1 and T2 samples. The random samples had higher alpha indices followed by T0, T1 and T2 samples. The Shannon alpha diversity index was found to be the lowest (3.45) in the faecal microbiome of the 10% maize feed diet, whereas the diversity as observed was highest (6.06) in the random diet.

**Table 2:** Species Diversity Observed in the Sample.

| SI No. | Sample Name | Observed OTUs | Shannon alpha diversity |
| --- | --- | --- | --- |
| 1 | R | 548 | 6.06 |
| 2 | T0 | 450 | 5.87 |
| 3 | T1 | 479 | 5.12 |
| 4 | T2 | 203 | 3.45 |

### Rarefaction curve

The rarefaction curve is a plot of the number of species as a function of the number of samples. The rarefaction curves were built using re-sampling of R, T0, T1 and T2 samples several times and then plotting the average number of species found on each sample. **Figure 6** illustrates the rarefaction curves for the faecal microbial population of the experimental animals. The lines in the curve represent the observed OTUs at 3% dissimilarity. The curves infer greater bacterial diversity in the random group followed by T1, T0 and T2 groups. It is observed that pigs fed with a 10% maize diet had lesser bacterial diversity compared to pigs fed with a non-fibre diet.

**Figure 6:**
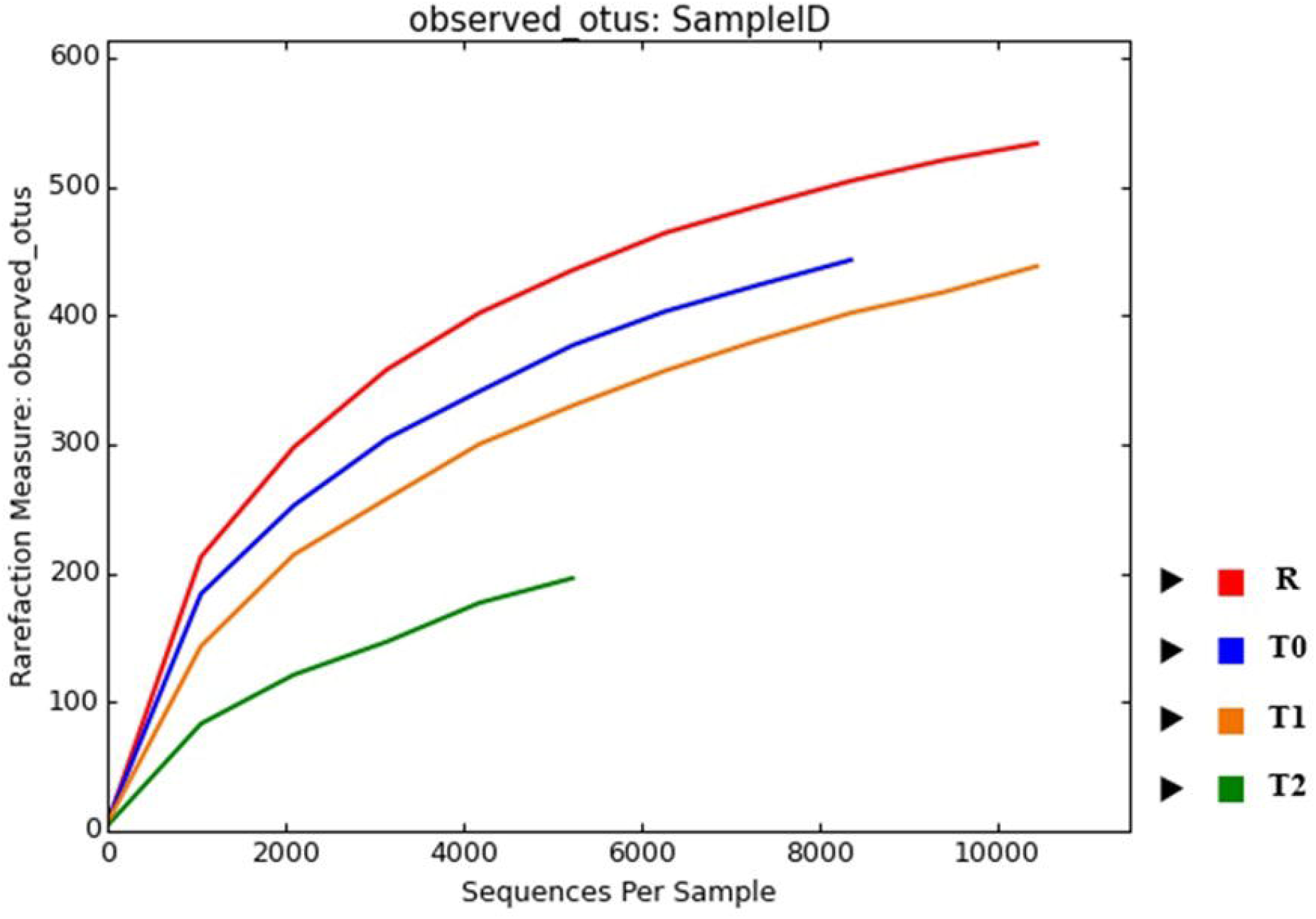
Rarefaction plot for samples R, T0, T1 and T2

### Heatmap

Heatmap images have been created to picture the OTUs at different taxonomic levels with each row corresponding to an OTU and each column corresponding to a sample (**Figure 7, Figure 8 and Figure S2**). The relatively higher abundance of an OTU in a sample was shown in the deeper colour at the corresponding position in the heatmap. The purple colour indicates a high percentage of OTUs to sample while the red colour indicates a low percentage. The abundance of phylum Fibrobacteres, was highest in the T2 group followed by T1 whereas it remained undetected in T0 and random. Analogously, the population of Spirochaetes increased from random to treated T2 groups. In contrast, the percentage of phylum Firmicutes, Bacteroidetes and Proteobacteria decreased with the increase in fibre content in the diet. The members of class Spirochaetes and Fibrobacteria was present at maximum in the T2 samples. The abundance of class Clostridia and Bacteroidea was highest in random and lowest in the T2 groups. Class Gammaproteobacteria and Methanobacteria were found to be highest in random samples. The profusion of Epsilonproteobacteria was large in T0 and T1 compared to T2, while it was negligible in the random. A similar drift in the bacterial microbiome was observed in the order, family and genus of the respective classes in each experimental group.

**Figure 7:**
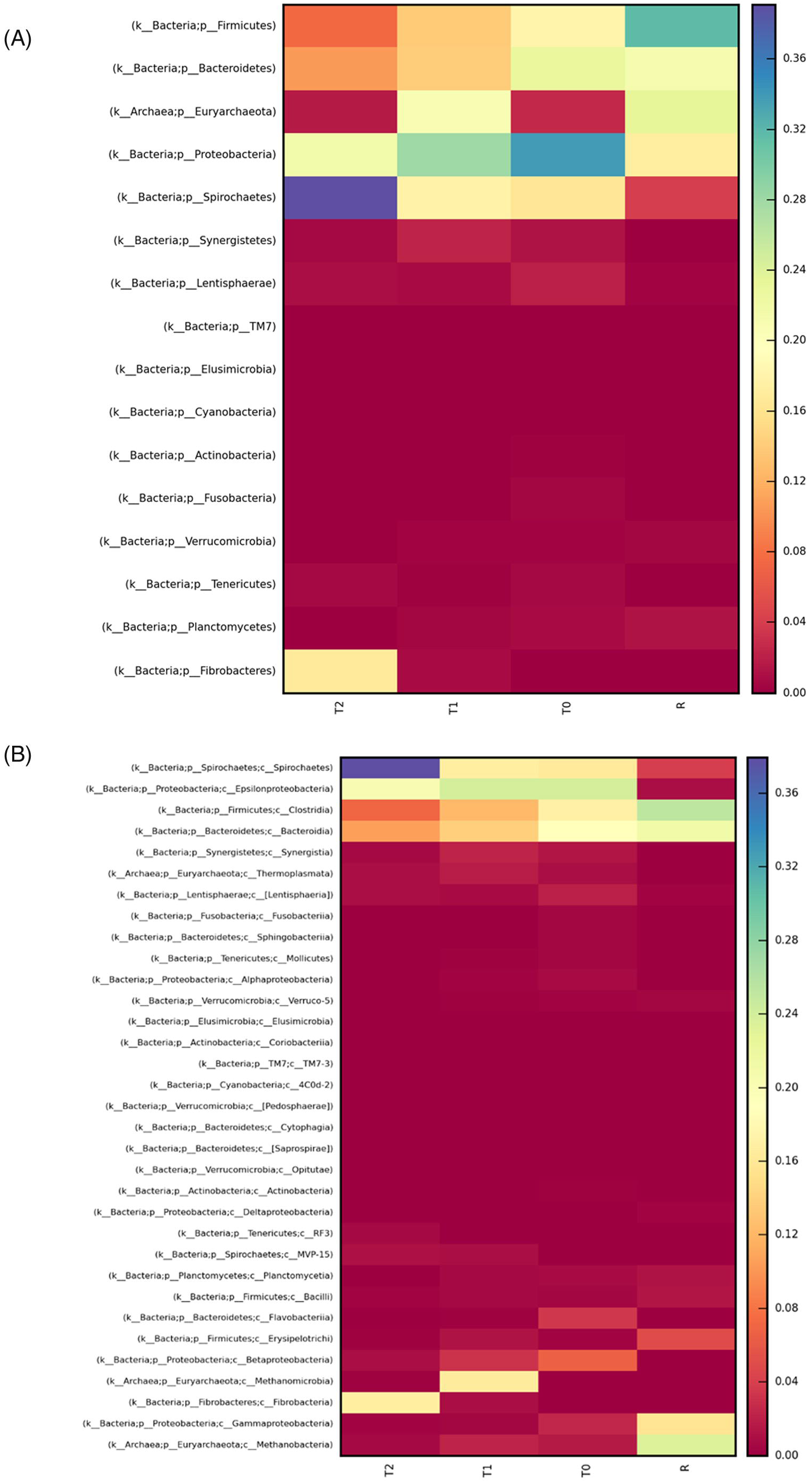
Heatmap visualizes the OTU table at **A**. Phylum and **B**. Class Level. The purple colour indicates a high percentage of OTUs to the sample and the red colour indicates a low percentage of OTUs.

**Figure 8:**
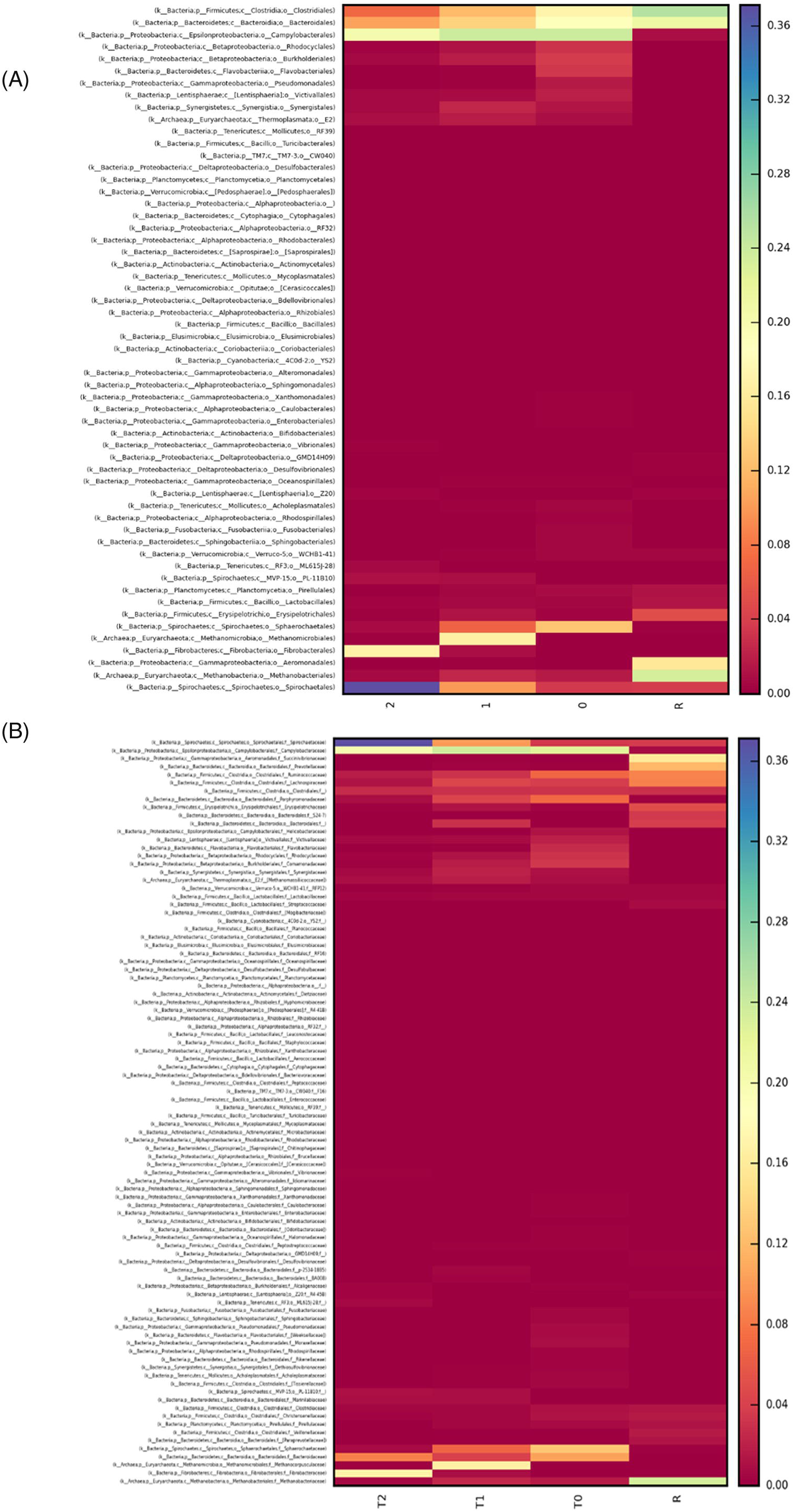
Heatmap visualizes the OTU table at **A**. Order, and **B**. Family Level.The purple colour indicates a high percentage of OTUs to the sample and the red colour indicates a low percentage of OTUs.

### Krona Charts

Krona plots are multilayered pie charts where taxonomic data can be viewed at multiple levels interactively. **Figure 9A, B, C** and **D** illustrate the multi-level Krona plot view of the bacterial community in R, T0, T1 and T2 samples. In the Krona chart, a whole metagenome is shown as nested concentric rings forming a circle together. Each of the rings indicates a single taxonomic hierarchy, the more distant the ring, the lower the strata. At each rank, a taxon is represented as a part of the ring that is equivalent to the abundance of the taxon in the sample.

**Figure 9:**
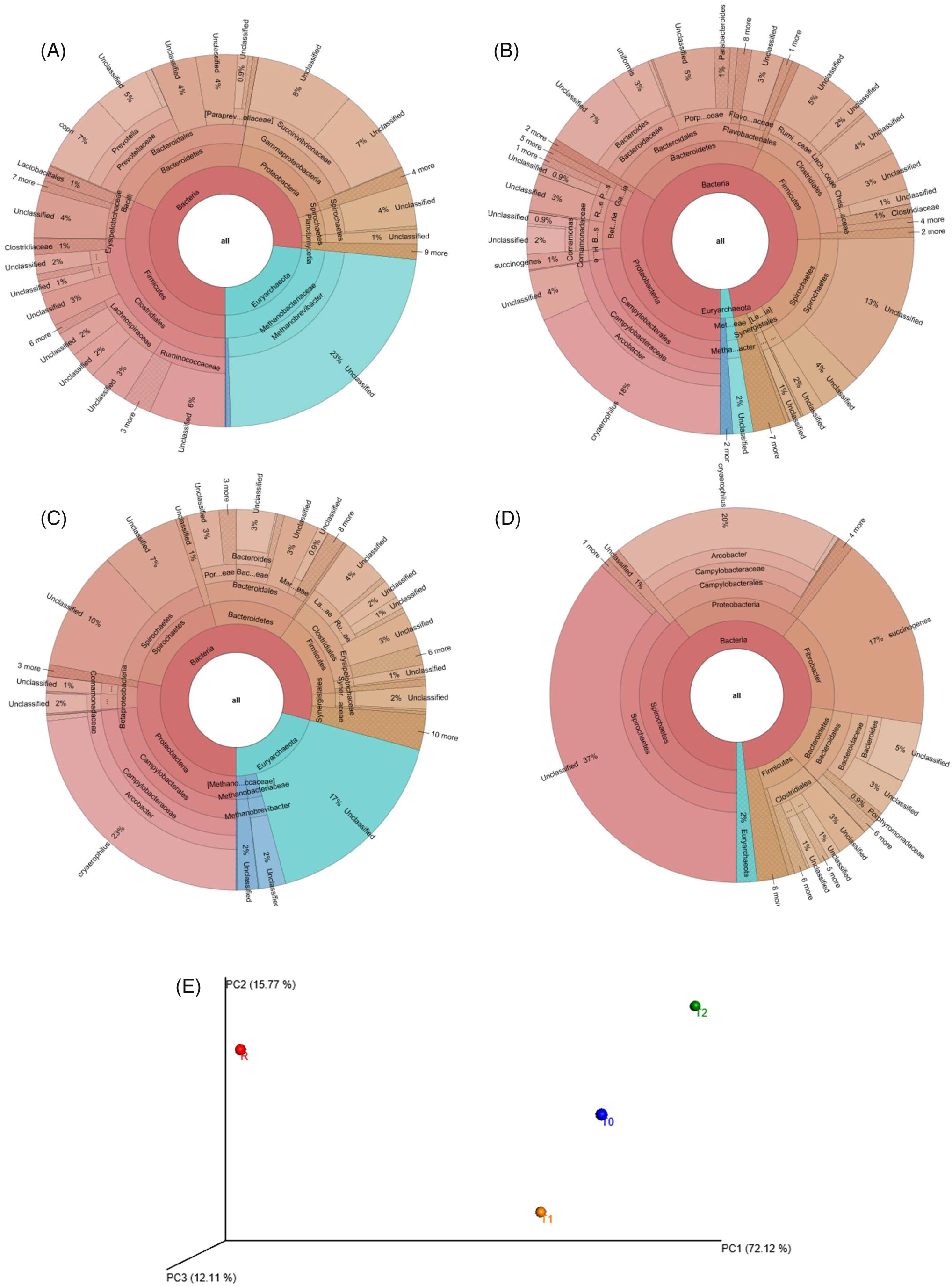
Krona chart showing taxonomic data at multiple levels in an interactive wayin **A**. R, **B**.T0, **C**.T1 and **D**.T2 samples and **E**. Principal coordinates plotted to provide a perceptive visualization of the data structure showing differences between the samples similarities by sample-based on weighted Unifrac distances of all samples (R-Red, T0-Blue, T1-Brown and T2-Green)

### Principal coordinate analysis (PCoA)

The beta diversity was estimated by QIIME software (version 1.8.0) with PCoA based on weighted Unifrac distances. **Figure 9E** shows the Principal Coordinate Plots of the experimental samples. The PCoA revealed that the random sample showed a more distant separation from the other three groups, while samples of T0, T1 and T2 were more similar to each other. Samples belonging to the random group were assembled far away from the other three groups which indicated that the gut microbiota of the random diet-fed pigs was strikingly different relative to the other groups. Though the samples of the group T0, T1 and T2 were close to each other, samples of T1 and T2 were closer to one another, signifying that these two groups of pigs have a similar kind of gut bacterial microbiome compared to the rest.

## DISCUSSION

A reference module of the swine gut microbiome was detailed in 2016^15^ by employing the metagenome sequencing records of 287 pig faecal samples collected from different countries. Diet is considered the indispensable factor in modifying the layout and functions of swine microbiota, which overcomes the differences in the host’s genetic parameter in determining the composition of the microbial community^29,30^. Previous reports have established the symbiotic association between gut microbiota and pig feed regulation^31,32^, growth^21^ and disease resistance in early-weaned piglets^15^. Breed precise bacteria in the swine intestinal chamber was reported to subsist, even though pigs were trialled with the same feed, farm ambience, and management measures^33^. A potential linkage between the microbiota intestinal and feed efficiency in pigs was deduced by Tan *et al*.^34^ stated that Landrace pigs with a high-feed conversion ratio had a greater accumulation of *Lactobacillus* and *Streptococcus* than the pigs with a low-feed conversion ratio. A greater profusion of *Oscillibacter, Christensenellaceae* and *Cellulosilyticum* was identified in the chamber of gastrointestinal track of Large White X Landrace pigs with high-feed efficiency^35^. Yang *et al*.^36^ pinpointed thirty-one OTUs in Duroc pigs addressing potential correlation with feed efficiency. Some previous studies reveal that in different varieties of pigs; there may be alterations in the composition of the microbial population with different advantageous species. The annotation alignment at the genera strata of the swine faecal microbial population interpreted different classification compositions. According to Yang and group^36^, *Lactobacillus*, *Prevotella*, and *Treponema* were the three most copious genera in Duroc pigs. On a similar note, *Prevotella, SMB53*, and *Streptococcus* were found to be the most abundant genera in Hampshire pigs, while in Landrace and Yorkshire, the three amplest genera were noted to be *Clostridium, Streptococcus* and *SMB53*^33^.

The current investigation underscored the conspicuous dominance of phylum Firmicutes, Bacteroidetes, Proteobacteria and Spirocaetes in every experimental group in different fractions. But the trajectory ratio of phylum Firmicutes, Bacteroidetes and Proteobacteria decreased with the increase in fibre content in the diet. Firmicutes, Bacteroidetes and Spirocaetes are the most abundant gut phylum reported in pigs^37^. Bacteroidetes are notably considered experts for the splitting of high molecular weight biomacromolecules like proteins, carbohydrates and organic matter^38^. They contain specific chemistry in chopping complex dietary polysaccharides into small-chain fatty acids, which can be readily reabsorbed by the host gut. Gut Bacteroidetes generally have antineoplastic properties and benefit the host’s health, by uniting with the immune orchestra to stimulate T cell-mediated responses^39,40^. Relative abundance of phylum Firmicutes was found to be associated with energy uptake from the feed content^41^. A higher quantity of Firmicutes was observed in obese animals and adults compared to lean individuals. Proteobacteria are known to occupy a unique space in the biome of microbiota since they contribute largely towards carbohydrate and protein metabolism and also maintain oxygen homeostasis in the gastrointestinal milieu of healthy mammals^42^. However, members of Proteobacteria, which comprises a wide diversity of pathogens (such as *Salmonella, Escherichia, Helicobacter, Yersinia* and *Vibrio*) are well recognized for their association with different diseases. An increase in the percentage of Proteobacteria has been predicted as a sign of dysbiosis by a wide group of studies^43–45^. According to Shin *et al^43^* Proteobacteria has also been found to be interlinked with obesity.

A striking observance of the current study is the subsiding abundance of Eurybacteria from the R to the T2 groups. Euryarchaeota is known as a prolific colonizer in the gastrointestinal route of certain mammals, especially herbivorous^46^. Euryarchaeota includes many methanogens that have the unique capability to produce methane through cellulose metabolism in hypoxic conditions^47^. However, methanogens found to be associated with dysbiosis such as in the case of vaginosis^48^, urinary tract infections^49^, and anaerobe abscesses of the brain^50,51^, the muscle^52^, the oral cavity in the case of periodontitis, and periimplantitis,^53,54^ in the case of refractory sinusitis^55^. In humans, intestinal methanogens are interrelated with numerous ailments such as irritable bowel syndrome, inflammatory bowel disease, obesity, constipation colorectal cancer, anorexia, periodontitis, diverticulosis, and colorectal cancer^56,57,46^. While, from the environmental outlook, methane production in gut fermentation is of global apprehension for its impact on the accretion of greenhouse gases in the atmosphere, as well as its waste of fed energy for the animal. Methane is synthesized in the intestine of animals by a section of Archaea, which belongs to the phylum Euryarcheota. Many scientific works have been focused on methane attenuation strategies to be used in animal husbandry^58–60^. Methane is synthesized in the gastrointestinal tract as a composite of the normal fermentation of feedstuffs. The composition of the diet supplied, specifically the type of carbohydrate consumed, is exclusive for methane building as they are responsible to influence the intestinal pH and consequently amend the present microbiota^61^. The digestibility of cellulose and hemicellulose are very much correlated with methane accumulation than soluble carbohydrates^62^. For decades, scientific communities have been contemplating identifying, enumerating, and annihilating methanogens and methanogenesis through different methane alleviation policies. The decrease in class methane bacteria in the fibre-treated pigs in the present study shows an obvious decrease in the production of methane and its diaspora in the environment and ultimately leads to healthy gut digestibility in the pigs.

The intestinal environment comprises diverse microbial accumulate that are specific to the different sections of its tract. The microbiome ensemble is markedly segregated among the small intestine, colon, and faeces, and the bacterial contour of the colon resembles much of that of faeces^63^. The small bowel is evidenced to have a lesser diversity and abundance of microbial corpus compared to the colon^64^. Swine faecal microbiota is particularly analogous to the colon microbiota, where digestion of complex carbohydrates takes place but is distinct extensively from the ileal microbiome which are experts on the anaerobic breakdown of mono- and disaccharides^65,66^.

The investigation highlights the increase in the colonization of the beneficial bacterial phylum Fibrobacteres with the increase in the percentage of green maize (T1 and T2), whereas no bacterial ASV of the said phylum was detected in the trialled pigs of low fibre diet (R and T0). The alpha diversity index was found to be low in the fibre-fed diet compared to the random diet. The random samples had higher alpha indices followed by T0, T1 and T2 samples. The alpha diversity of the gut microbiota as per our findings did not increase concurrently with the change in dietary supplements. The findings of this study parallel with those of Li *et al*.^67^, which showed the alpha diversity (Shannon index and observed OTUs) of diet was not concomitant with that of swine gut microbiota. The multifariousness of the gut microbiome was not linearly correlated with the increase in the diversity of foods. In one of the published data Bolnick *et al*.^68^, have validated that different feeds have non-additive effects on microbial contexture in the gut of fish^68^. A similar trend of bacterial species richness was figured out from the rarefaction analysis, which showed the highest diversity in random diet and lowest in fibre diet. As reported earlier^67^, some microbes act as dominants after they have attained competitive advantages in the gut as they have a broad variety of nutrients in the intestinal surroundings. While on the other half, some rare microbes having a narrow zone of nutrients fail to survive in the host atmosphere. Additionally, every specific diet might prompt a chemical inhibitor that stimulates the existence or proliferation of specific microbes. As an outcome, dietary diversity may not increase in lockstep with gut microbial diversity. As shown in this study, the diet-induced shifts in beneficial microbial bloom may influence overall fibrolytic activity in swine guts.

Genus *Fibrobacter* (which increased after feeding on green maize) is generally regarded as an efficient and unique bacterial degrader of substances like lingocellulosic forages in the gastrointestinal tract^69^ and is extremely copious in animals served with low-quality nutrients. It has the efficiency to break down crystalline cellulose and solubilize plant polysaccharides^70^. Equally, the genus *Treponema*^71^ is also known to be involved in fibre degradation. Li *et al*.^67^ recommended that individuals that consume an analogous diet harbour more analogous gut microbiota in their intestinal ambience. The results of the present study were corroborated with earlier findings of forage effects over rumen microbial contexture in dairy cows^72^. As studied by Xie and his group^73^, feeding of corn stover ascertained the proliferation of lignocellulose chopper in the rumen of Hu sheep and transformed the entire microbiome towards a better fibrolytic purging ecosystem in comparison to alfalfa hay. Ingestion of natural high lignocellulose feedstuffs for a longer period facilitates higher fibre digestion capacity in wild ruminants such as bison, buffalo, and yak compared to domestic bovines^74^. Even monogastric animals showed persistent gastrointestinal microbiome patterns, which were formed under long-term dietary regimes^75^. Feed intervention can switch to microbiome modification while keeping the differences caused by individuality and host resilience sideways^73^. Liu *et al*.,^76^ found that cereal straw can act as a regulator and enhance the function of fibrolytic bacteria in the gut of ruminants. Even though microbes’ have the speciality to rapidly acclimatize to a new diet, some microbial members can remember the former dietary experience and persevere under the new diet, as demonstrated in monogastric animals^77^. Parallel with previous findings in ruminants fed with a forage-based diet^78^, we identified relatively higher abundances of *Fibrobacter* and *Treponema*, in fibre-fed samples in the present study, signifying that these taxa enact essential roles in occupying specific niches in pig gut ecology and the degradation of forage-based feedstuffs. *Fibrobacter* and *Treponema* have been shown to have a mutual interrelation^79–81^, in which *Fibrobacter* counts on *Treponema* to splice out interlaced hemicellulose to gain easy access to fibre components, and *Treponema* in turn relies on *Fibrobacter* to provide cellulose hydrolysates. Different foodstuff indirectly affects not only host physiology but also affects the immunity of the host, which may regulate the diversity of gut microbiota according^67^. On the other hand, a lack of alpha diversity indicates that there have been positive correlations among gut microbiota, beta diversity and diet.

## CONCLUSION

The present investigation reports a dynamic microbial architecture in the faecal microbiota of pigs fed with different levels of maize fodder. The study has detected a lower bacterial diversity in fibre-fed T2 samples with 203 OTUs compared to the low-fibre-fed samples with 548 OTUs. The microbiome of the T2 groups was concentrated essentially of the phylum Firmicutes, Bacteroidetes, Proteobacteria, Spirocaetesand Fibrobacteres. Colonization of phylum Fibrobacteres was detected only in the fibre-fed diets underscoring that feeding of green maize has stimulated the increase of fibrolytic bacterial genera- *Fibrobacter* and *Treponema* compared to the low-fibre diet. The synergistic effect of *Fibrobacter* and *Treponema* are found to have a superior aptitude to degrade dietary cellulose, polysaccharide and protein, encouraging good intestinal health. The hypothesis concludes that feeding of fibre enriched diet has acclimatized the persistence of fibrolytic bacteria- *Fibrobacter* and *Treponema*, which has further increased the fibre digestibility in the monogastric animals. Thus, feeding with a fibre-enriched diet has augmented healthy gut microbiota in pigs and led to the dominance of beneficial microbes that are quintessential to enhancing the gastrointestinal milieu of the porcine animals. Nevertheless, feeding green maize has decreased the population of methane bacteria in the gut of pigs, which in turn has limited the production of methane. A fibre-rich diet is a good remedy for improving intestine digestibility in pigs, and it can also be used as a strategy to reduce global warming’s impact.

## MATERIALS AND METHODS

### Statement

The study was conducted at the Indian Council of Agricultural Research-National Research Centre on Pig, Rani, Guwahati, 781131, India. The experimentation was carried out with prior approval from the Institute Animal Ethics Committee. The approved animal use protocol number is NRCP/CPCSEA/1658/IAEC-52 dated 03-12-2019. All experiments were performed in accordance with relevant guidelines and regulations.

### Design of Experiment and collection of sample

A total of 24 grower pigs of the same age, gender, breed, weight and same genetic backgrounds were selected and raised with a similar customized diet. The experimental pigs aged three months were reared in the controlled farm conditions with uniform management conditions and divided into four groups T0, T1, T2 and R. The average body weights of the experimental animals were 30.67±1.84, 30.47±1.04, 30.04±0.76 and 31.04±0.36 kg respectively in R, T_0_, T_1_ and T_2_ group. The experiment was conducted for three months. The lysine and methionine were balanced in all the rations as per requirement. The calculated energy^24^ (ME, Kcal/kg) of the experimental diet was 3345, 3327.9 and 3321.8 respectively in T_0_, T_1_ and T_2_ groups and the energy (ME, Kcal/kg) value in the R group was 3365.2.

Experimental animals were supplemented with 0, 5 and 10 % QPM maize fodder to the basal diet by replacing (wt/wt on DM) the maize grains and designated as T_0_, T_1_ and T_2_ respectively. R group was fed with a normal diet comprising maize crush-60 kg, wheat bran: 11 kg, ground nut cake: 13 kg, soybean meal: 13.5 kg; mineral mixture 2kg and common salt 0.5 kg. All the experimental diets were iso-nitrogenous containing 18% protein. Dry matter intake was found similar across the treatment groups ranging from 1313 to 1319 g/d respectively in T_2_ to T_0_ and others were within the range of variation. There was no significant difference in nutrient digestibility across the treatment groups. Similarly, there was no significant difference in average daily gain (g/day), feed intake per kg gain (FCR) and feed cost per kg gain. However, FCR and feed cost per kg gain were found better at 5 % and 10 % supplementation of maize fodder in the diet. Average daily gain (g/day) ranged from 304.77 to 306.25 respectively in T_2_ and T_0_ groups.

The metagenomic profiles of pig gut microbiota of four different samples (random, green maize 0%, 5% and 10%) have been analyzed. Pig faecal samples of four different categories *viz*. R (Random), T_0_, T_1_ and T_2_ animals were collected after stimulation of the rectum and samples were transferred quickly to liquid nitrogen for temporary storage. Subsequently, the samples were stored at −80°C in the laboratory until analysis. DNAs were isolated from 24 numbers of faecal samples from all the four categories (R, T_0_, T_1_ and T_2_) and then pooled the DNA of each group containing six numbers animals, which was further used for metagenomic sequencing. A comprehensive comparative analysis of these samples was conducted at phylum, class, order, family, genus and species levels to identify the beneficial bacterium responsible for fibre degradation as well as to find out the most abundant bacteria present in the pig gut.

### Metagenomic DNA Extraction, Qualitative and Quantitative analysis

The commercially available QIAamp DNA Stool Mini Kit (Qiagen Ltd., Germany) was used to extract DNA from the pig faecal samples following the manufacturer’s protocol. The concentration of DNA was measured using a spectrophotometer, and 1% agarose gel electrophoresis was run to check the integrity of the DNA of the faecal sample. Finally, a 260/280 ratio of 1.75-1.85 of DNA samples was considered for the downstream analysis. TheV3 and V4 regions of bacterial 16S rRNA genes were amplified from pig faecal samples DNA using Taq polymerase with primers (F: 5’-GCCTACGGGNGGCWGCAG-3’ and R: 5’-ACTACHVGGGTATCTAATCC –3’).

### Preparation of 2 x 300 MiSeq libraries

The Nextera XT Index Kit (Illumina inc.) was used to construct the libraries. The metagenomic libraries for DNA of pig faecal samples were constructed with an insert size of 350 base pairs (bp) for each sample using Illumina protocol^25^ and the PE150bp strategy were used to sequence the 16S Metagenomic Sequencing Library following protocol (Part # 15044223 Rev. B) of MiSeq 2500 platform (Illumina). The amplified 16S gene by PCR was resolved in a 1.5% agarose gel.

### Check of Quality and quantity of the library

The Agilent 4200 Tape Station was used to amplify library sequences and was using D1000 Screen tape as per manufacturer protocol.

### Quality Control (QC)

The i5 and i7 primers were used to amplify the Illumina adaptor that has passed the QC and subsequently multiplexing was added to index sequences as well as common adapters sequenced that were required for cluster generation (P5 and P7) as per the standard manufacture protocol. The low qualities of raw reads were either trimmed or removed for further analyses, as well as reads with an adaptor or more than three nitrogen bases, were removed. Further 30 ends of the read were trimmed and bases with low quality (Phred score < 20) were removed for further analyses if the original read length is shorter than 50 bp. Subsequently, the SOAP aligner (v 2.21)^26^ was used to remove host (pig) genomic DNA sequences. The Qubit Fluorometer was used to quantify the amplicon libraries purified by AMPure using XP beads strategies.

### Generation of Cluster and Sequencing

The concentration of 10-20 pM was used as a benchmark for cluster generation and sequencing after obtaining the mean peak sizes from the Agilent Tape Station profile, subsequently, libraries were loaded onto MiSeq at an appropriate concentration for cluster generation and sequencing. Paired-End sequencing allows the template fragments to be sequenced in both the forward and reverse directions on MiSeq. All the samples were bound to complementary adapter oligos on the paired-end flow cell using specific kit reagents. During sequencing, selective cleavage was produced in forwarding strands after re-synthesis of the reverse strand by the designed adapters. Subsequently, sequencing was done from the opposite end of the fragment of the copied reverse stand.

The Nextera XT Index Kit (Illumina Inc.) was used for the first amplicon generation followed by NGS library preparation of the QC passed samples. The QC passed libraries were sequenced on the MiSeq platform (Illumina) using 2 x 300bp chemistry to generate ~1 lakh reads per sample.

### Bioinformatics Analysis

The QIIME (version 1.8.0) bioinformatics pipeline is used to analyse the faecal microbiome from raw DNA sequencing data. This bioinformatics pipeline is very strong and comprehensive that comprises heuristic-based maximum-likelihood phylogeny inference that generates FastTree^27^ as well as also used to assign taxonomic data through naive Bayesian classifier^28^. Trimmomaticv 0.38 software was used to identify high-quality clean reads. This software specifically removes the adapter sequences, ambiguous reads (“N” > 5%), and low-quality sequences (reads > 10% quality threshold (QV) < 20 Phred score) along with a sliding window of 10bp and a minimum length of 100bp. Finally, all the Paired-End (PE) sequence data were aligned into single-end reads.

The representative sequence for 16S Bacteria DNA was retrieved from each Operational Taxonomic Unit (OTU) against the Green genes database (version 13_8). The OTUs are based on sequence similarity within the reads and pick. Subsequently using reference databases, OTUs were assigned a taxonomic identity and calculated diversity metrics. Finally, a PCoA (Principal coordinate analysis) plot was created visually to illustrate the differences between the samples.

The OTUs of highly similar sequences across the sample were pickup to provide a platform for comparisons of community structure as well as showing some degree of taxonomic relatedness by using UCLUST at 97% sequence similarity. This has resulted in the formation of a cluster that typically represented a species. For downstream analysis, only one representative sequence of OTU is picked out of many sequences.

## ACKNOWLEDGMENTS

The authors are grateful to the Director, ICAR-Indian Institute of Maize Research, PAU Campus, Ludhiana 141 004, India for financial assistance under the NEH grant of the Institute. The authors are also grateful to the Director, ICAR-National Research Centre on Pig, Rani, Guwahati, 781131, India for providing the facility to conduct the experiment.

## AUTHOR CONTRIBUTIONS

KB, PD, SJ, SR and VG conceived the idea and designed the experiments, and edited the manuscript. MC KB, SR, SB and PD analyzed the result and draft the manuscript. SP, SK, RD and JR performed the collected samples experiments and run the experiments. All the authors reviewed and approved the manuscript.

## DATA AVAILABILITY STATEMENT

The compete metagenomic data sequenced on the Illumina MiSeq platform were deposited in the NCBI’s Sequence Read Archive (SRA) database under the accession ID PRJNA795318.

## ETHICS STATEMENT

The study was conducted at the Indian Council of Agricultural Research-National Research Centre on Pig, Rani, Guwahati, 781131, India. The experimentation was carried out with prior approval from the Institute Animal Ethics Committee. The approved animal use protocol number is NRCP/CPCSEA/1658/IAEC-52 dated 03-12-2019.

## SUPPLEMENTARY MATERIAL

The Supplementary Material for this article is mentioned below.

**Figure S1**: Bar chart (Stacked) showing the relative abundance of each **A**. Genus and **B**. Species within each sample

**Figure S2**: Heatmap visualizes the OTU table at Genus Level. The purple colour indicates a high percentage of OTUs and the red colour indicates a low percentage of OTUs.

