## Supplementary figures and images for "Metagenomic analysis of Pigs’ faecal microbiome and its functional response associated with dietary fibre"

### Figure S1

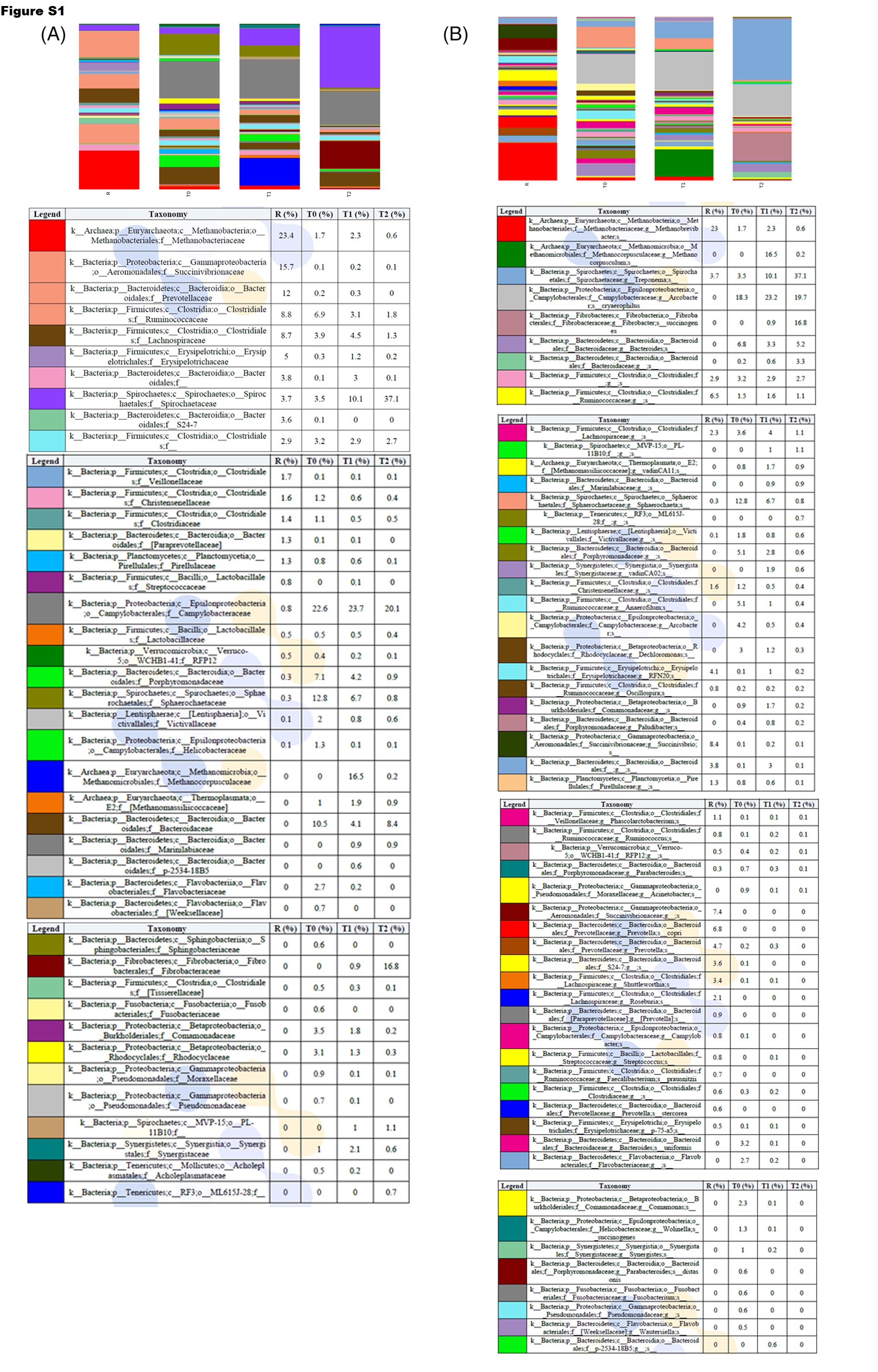

### Figure S2

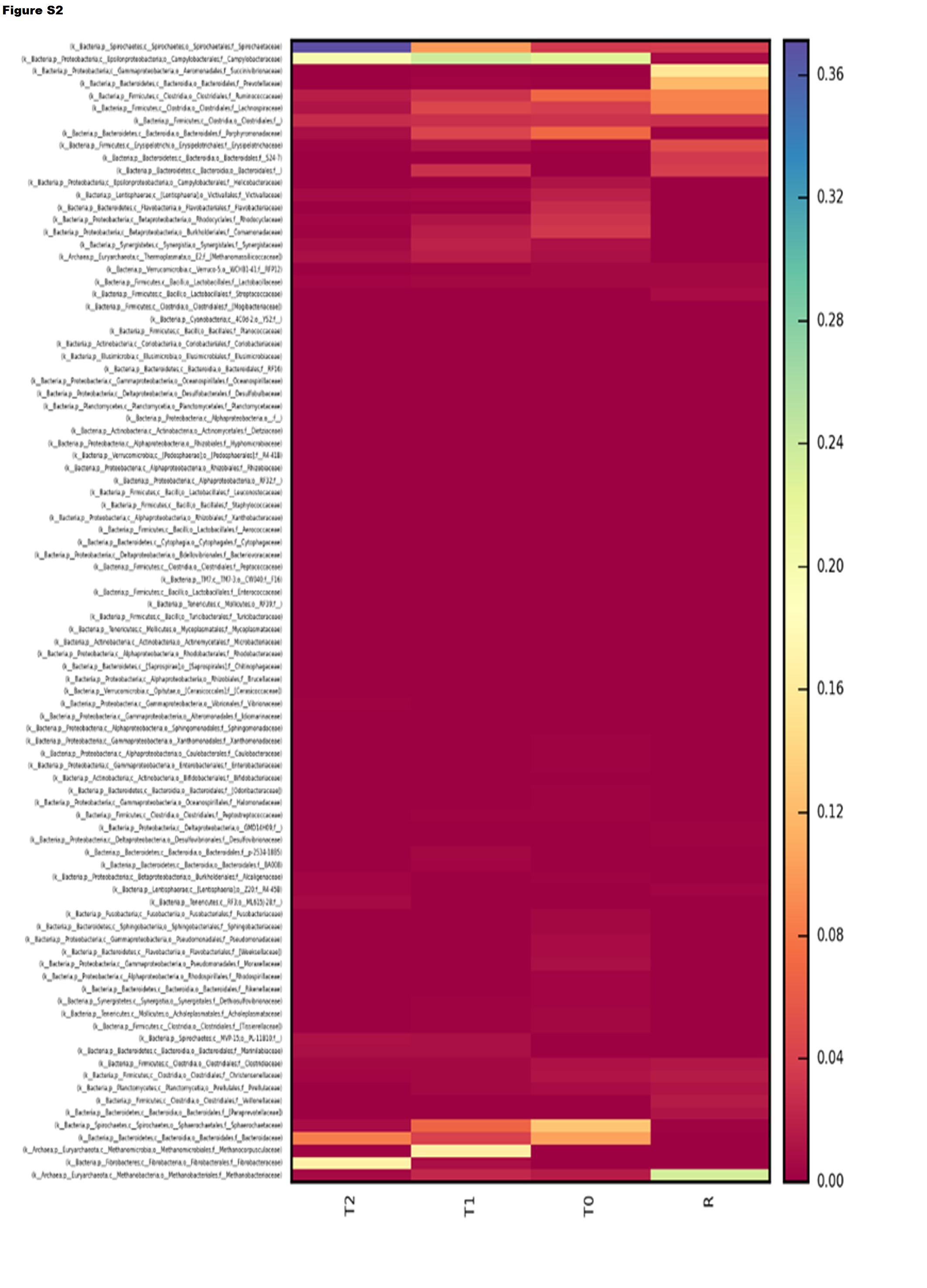
